# LIONirs: flexible Matlab toolbox for fNIRS data analysis

**DOI:** 10.1101/2020.09.11.257634

**Authors:** Julie Tremblay, Eduardo Martínez-Montes, Alejandra Hüsser, Laura Caron-Desrochers, Philippe Pouliot, Phetsamone Vannasing, Anne Gallagher

## Abstract

**Background:** Functional near-infrared spectroscopy (fNIRS) is a suitable tool for recording brain function in pediatric or challenging populations. As with other neuroimaging techniques, the scientific community is engaged in an evolving debate regarding the most adequate methods for performing fNIRS data analyses.

**New method:** We introduce LIONirs, a neuroinformatics toolbox for fNIRS data analysis, designed to follow two main goals: (1) flexibility, to explore several methods in parallel and verify results using 3D visualization; (2) simplicity, to apply a defined processing pipeline to a large dataset of subjects by using the MATLAB Batch System.

**Results:** Within the graphical user interfaces (DisplayGUI), the user can reject noisy intervals and correct artifacts, while visualizing the topographical projection of the data onto the 3D head representation. Data decomposition methods are available for the identification of relevant signatures, such as brain responses or artifacts. Multimodal data recorded simultaneously to fNIRS, such as physiology, electroencephalography or audio-video, can be visualized using the DisplayGUI. The toolbox includes several functions that allow one to read, preprocess, and analyze fNIRS data, including task-based and functional connectivity measures.

**Comparison with existing methods:** Several good neuroinformatics tools for fNIRS data analysis are currently available. None of them emphasize multimodal visualization of the data throughout the preprocessing steps and multidimensional decomposition, which are essential for understanding challenging data. Furthermore, LIONirs provides compatibility and complementarity with other existing tools by supporting common data format.

**Conclusions:** LIONirs offers a flexible platform for basic and advanced fNIRS data analysis, shown through real experimental examples.

**Highlights:**

- The LIONirs toolbox is designed for fNIRS data inspection and visualization.
- Methods are integrated for isolation of relevant activity and correction of artifacts.
- Multimodal auxiliary, EEG or audio-video are visualized alongside the fNIRS data.
- Task-based and functional connectivity measure analysis tools are available.
- The code structure allows to automated and standardized analysis of large data set.

**Graphical abstract:** 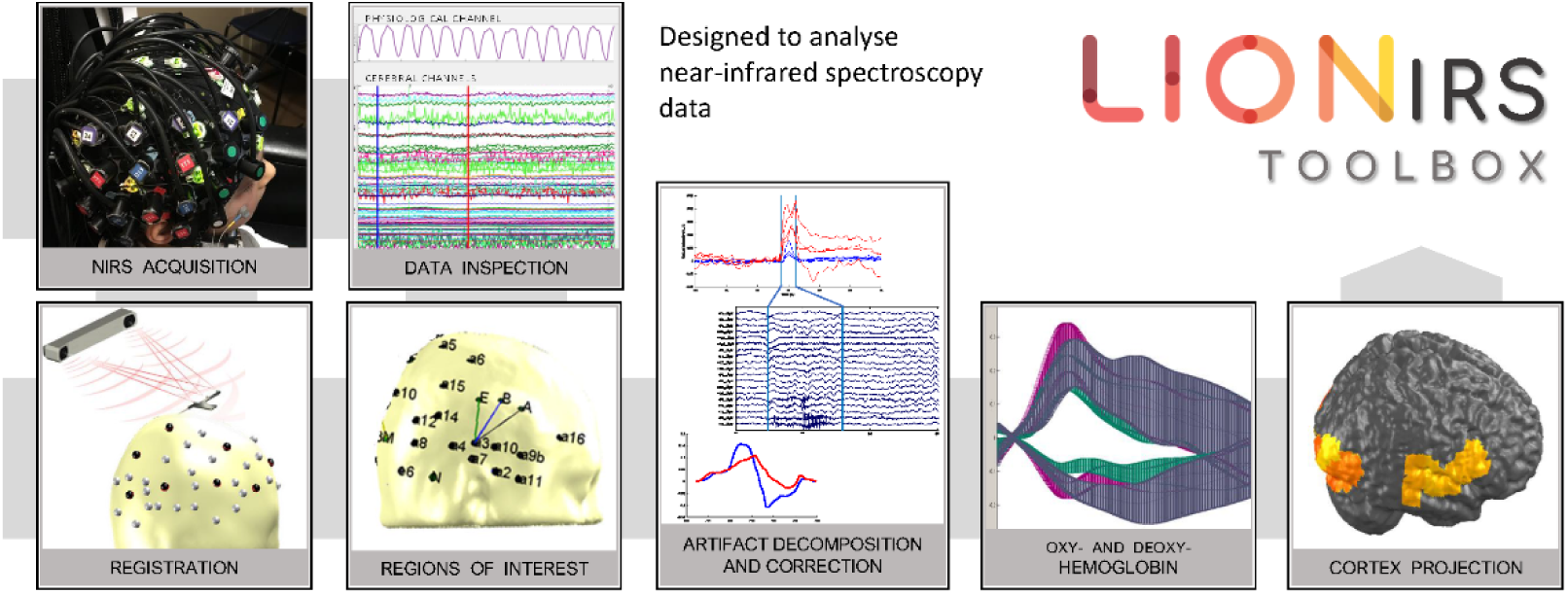

## 1. Introduction

Functional near-infrared spectroscopy (fNIRS) is a non-invasive neuroimaging technique. Similar to functional magnetic resonance imaging (fMRI), it measures blood oxygenation fluctuations related to neuronal processes in cortical regions, but with several practical advantages. First, it is portable, allowing studies to be conducted in clinical settings where bedside acquisition is necessary (Kassab et al., 2018). Second, fNIRS is relatively tolerant to movement, which makes it suitable for language studies that require participants to speak aloud. Third, fNIRS is child-friendly because it allows the researcher or a parent to interact with the participant during data recording. Hence, it is a technique particularly useful for challenging populations, such as infants and toddlers, or individuals with cognitive or psychiatric conditions (Vannasing et al. 2016). Finally, fNIRS can easily be used in multimodal contexts, such as during simultaneous fNIRS-EEG acquisitions, which could, for example, contribute to a better understanding of neurovascular coupling and brain function in patients with epilepsy (Wallois et al., 2012, Wallois2010).

Since fNIRS is a relatively young modality, there exists a wide variety of approaches for the processing and analysis of its data, but no clear consensus on the best way to apply them. For instance, it is not yet clear which are the most appropriate and standardized pipelines through which to extract relevant information in specific experimental settings. Therefore, flexible neuroinformatics tools are needed, in order to allow the exploration of several options for the processing and handling of fNIRS data. It is important for such tools to provide user-friendly approaches for dealing with theoretical issues (e.g. which artifact-correction or data analysis method is the most appropriate), as well as practical concerns (e.g. definition of optode and data locations, visualization, multimodal integration, the handling of large datasets, performance of quality checks).

As with any other neuroimaging technique, a major challenge with fNIRS data analysis is the separation of the relevant signal from confounding signals (e.g. movement artifacts, physiological artifacts related to systemic fluctuations). Although a careful installation of the cap that holds the optodes and good cooperation from the participant during data acquisition can both significantly minimize artifacts, the fNIRS signal will always contain some confounding signals. For instance, in pediatric populations, young children or participants with severe cognitive or behavioral deficits may be unable to stay still and focused for long periods of time; although the experiment is often temporarily interrupted to allow the child to move and relax, a large amount of data can sometimes be significantly corrupted by movements. Should this occur, it is crucial to proceed to artifact detection, and to then either correct or reject the contaminated data segments (Di Lorenzo et al., 2019). Appropriate methods should be used to reliably and efficiently detect and/or correct artifacts resulting from the participant’s movements or physiology (e.g. heartbeat, respiration, Mayer waves). Compared to other neuroimaging techniques, fNIRS is still far from having standard data analysis procedures, and inadequate signal processing could lead to incorrect interpretation of the data and unreliable results (Brigadoi et al., 2014; Di Lorenzo et al., 2019).

Several methods have been suggested for dealing with motion or muscular artifacts in the fNIRS signal (Schecklmann et al., 2017), none being ideal and each having their own pros and cons (Hocke et al., 2018). Depending on the nature of the artifact itself, the mathematical assumptions made within a method for isolating the artifact may not be appropriate. Some methods reduce the interference caused by artifacts by using spline interpolation (Scholkmann et al., 2010), while others decompose the data using target principal component analysis (tPCA) and then remove the components related to the main artifacts (Yücel et al., 2014). More recently, parallel factor analysis (PARAFAC) (Hüsser et al., 2019) has been used for decomposition, as this method takes advantage of the multidimensional nature of fNIRS data (time, space and wavelength), and the fact that movement artifacts equally affect both wavelengths at the same location (Cui et al., 2010). Independent component analysis (ICA) has largely been used in EEG, and magnetoencephalography (MEG) and has proven to be efficient for removing eye blinks, which generate systematic and reproducible artifacts that can be considered statistically independent from brain activity (Hyvarinen, 1999). The technique of ICA has also been used to unmix the independent components in fMRI and fNIRS (Beckmann and Smith, 2004; Hiroyasu et al., 2013). The use of wavelet decomposition has been proposed for reducing spike artifacts, by removing high-frequency outliers from the data (Molavi and Dumont, 2012). Recently, a hybrid two-step approach combining spline interpolation and wavelet decomposition has been applied to fNIRS data to correct artifacts caused by jaw movements related to articulation during the performance of a language paradigm (Novi et al., 2020). Finally, a least-squares regression method has also been used to measure, estimate and subtract physiological artifacts caused by systemic fluctuations (Saager and Berger, 2005; Pfeifer et al., 2017). It is not yet clear which method is best for fNIRS artifact detection and correction, and the method chosen certainly depends on the type of artifacts being dealt with.

Furthermore, many approaches can be applied for the subsequent analysis of the artifact-free data. A task-based paradigm will assess the overlap between the recorded brain response and the hemodynamic response function (HRF), which is modeled using a general linear model (GLM) (Uga et al., 2014; Scholkmann et al., 2014; Peng et al., 2016). According to some studies, predefined theoretical HRF models can be problematic, notably in studies with clinical populations, infants or the elderly, or when focusing on complex cognitive functions (Nguyen et al., 2012, Nguyen et al., 2013, West et al., 2019, Buchsbaum et al., 2005). Moreover, there are some nonlinearity effects on the HRF response that are not yet completely understood (Zhang 2014). The higher temporal resolution at the acquisition of ∼0.05s for fNIRS in comparison to ∼1s for fMRI (Glover, 2011; Witt et al., 2016) could help describe the HRF nonlinearity and bring additional information through its time course morphology. Other avenues may explore functional connectivity measures to describe cognitive brain function at rest. Considering this large range of existing methods, one could easily feel lost, especially as few of the methods require advanced programming skills.

The development of more user-friendly tools, notably through graphical interfaces for the processing and visualization of data, is critical for applied and clinical research teams. These tools should offer several flexible methods and functionalities, allowing the user to create the best pipeline for the analysis of their data set. Since the analysis of a sizeable number of subjects may be extremely laborious, and even unrealistic in very large studies, some parts of the process are best automated, once the ideal pipeline has been created. However, as described above, a fully automated method for removing artifacts from fNIRS data may not be the best option, as inadequate use of signal decomposition poses the risk of distorting or even removing the signal of interest itself. It is therefore crucial that the user have access to various artifact detection/correction methods, to be able to identify the most appropriate one for a given data set. The chosen method would then be applied in a semi-automated manner to the entire sample of subjects. However, a supervised application is especially important for noisier data sets. The user should be able to assess the signal quality with a visual inspection before, during, and after the application of artifact detection and correction. The optimal tool would also allow the user to simultaneously access the data acquired from multiple modalities (physiology, EEG, audio-video recording, etc.), which has been shown to help with the identification of different events in fNIRS data (movements, epileptic events, state of consciousness, etc.) (Louis et al., 2016). Finally, in fNIRS, the placement of optodes is not standardized, but is instead customized in function of the cap used, the number of optodes available in the equipment, and the experimental question being posed. A tool that allows the projection of data onto either a brain or scalp reconstruction or a template can therefore greatly help with the visualization of data.

Several good neuroinformatics toolboxes have been developed to help researchers perform fNIRS data analysis. One of the first and best known is HomER, which is publicly available on the NITRC web site (Huppert et al., 2009). It is easy to use and offers a visual display of various options for motion correction and preprocessing. As of 2015, it also includes an external 3D interactive visualization of the data’s spatial distribution in AtlasViewer (Aasted et al., 2015). The Brain AnalyzIR toolbox is the most flexible tool available (Santona et al., 2018); it offers various statistical methods for addressing the specific features of fNIRS data analysis (e.g. specific models for the noise covariance). However, AnalyzIR requires relatively advanced programming skills, which are not typically available in applied and clinical research teams. NIRS-SPM (Ye et al., 2009) applies the statistical GLM method, which leads to the creation of activation maps, but it does not include modules for artifact rejection, which is critical, especially in studies with infants and children. Finally, FC-NIRS focuses mainly on the computation of functional connectivity matrices, based on temporal correlations of fNIRS time series (Xu et al., 2015).

In summary, there is currently no available toolbox that provides transparency and flexibility for data quality control, and a single solution for all the challenges faced when performing fNIRS data analyses, such as the need for an interactive 3D visualization adapted for a large number of channels, the flexibility to explore different methods for building customized processing pipelines, and offering multimodal integration so as to fully benefit from any additional source of information acquired simultaneously. These elements are all crucial for data sets acquired from challenging populations, such as young children or patients, where understanding the signal’s structure requires efficient high-density data visualization at each step of processing.

In this article, we introduce the LIONirs toolbox as a new tool for the analysis of fNIRS data. LIONirs is intended to complement other existing software, by supporting compatible file formats. It offers the flexibility to explore and visualize the data at each step of its processing, and allows one to easily build a customized processing pipeline that can be applied to a large number of subjects. For this purpose, the LIONirs toolbox is embedded in the Matlab Batch System (http://sourceforge.net/projects/matlabbatch/), used in the SPM toolbox (Penny et al. 2011). This allows the inclusion of various modules for data handling and processing. It also provides the ability to easily standardize the analysis of large data sets by the use of processing pipelines or templates. In this paper, the reader will be guided through the organization of the LIONirs toolbox, and its applications are illustrated with examples. The toolbox is available under the General Public Licence (GNU) and maintained on the public github.com project (https://github.com/JulieTremblay3/LIONirs).

## 2. Methods and results

### 2.1. Software overview

The LIONirs toolbox is an open-source software written entirely in Matlab (The MathWorks, Inc., Natick, Massachusetts, United States; compatible with releases from 2014a). It was developed in Windows, using the batch data editor in SPM12, and offers a variety of either basic or more sophisticated functions for handling and processing fNIRS data. It is therefore a minimal requirement that the user install the Statistics and Machine Learning Toolbox and SPM12.

An important characteristic of the LIONirs toolbox is its modularity. It offers the user the possibility of performing data processing in a customized manner and to integrate all steps into a processing pipeline. The pipeline can be saved as a template to be used for subsequent analyses. The user can also prepare a hierarchical pipeline by creating different branches for testing alternative processing approaches. Branches could, for instance, correspond to two different conditions (e.g. resting state during different stages of alertness: sleep vs. awake) or to different types of artifact correction (e.g. PCA vs. PARAFAC). The corresponding Matlab structure for each branch is saved in separate subfolders, allowing the user to easily assess the results of each individual processing approach. All these operations and their corresponding parameters are codified and stored in a Matlab structure named NIRS.mat.

The main graphical interface in the toolbox is called DisplayGUI. This interface uses the NIRS.mat structure to help the user modify artifact selection, and to easily and simultaneously visualize fNIRS data and multimodal data (e.g. heartbeat, respiration, EEG, audio-video recordings), using topographical projections onto the skin or cortical surface. Moreover, automatic script generation is available for creating a template, which can then be applied for multiple subjects [https://sourceforge.net/projects/matlabbatch/].

Figure 1 gives an overview of the organization and functionalities of the LIONirs toolbox. One of the first steps in fNIRS data analyses is to provide a 3D representation of the association between the positions of the optodes and the anatomy of reference. The toolbox includes the 3DMTG graphical interface, which allows the user to associate the raw fNIRS data with the corresponding 3D representation of the montage and the anatomy. First and foremost, LIONirs offers a flexible and semi-automated artifact correction strategy by use of various data decomposition methods, such as tPCA or PARAFAC. Once the signal is considered clean, or without artifacts, transformation of the light intensity into concentrations of oxy- and deoxyhemoglobin is done using the Modified Beer-Lambert law (MBLL).

**Figure 1.**
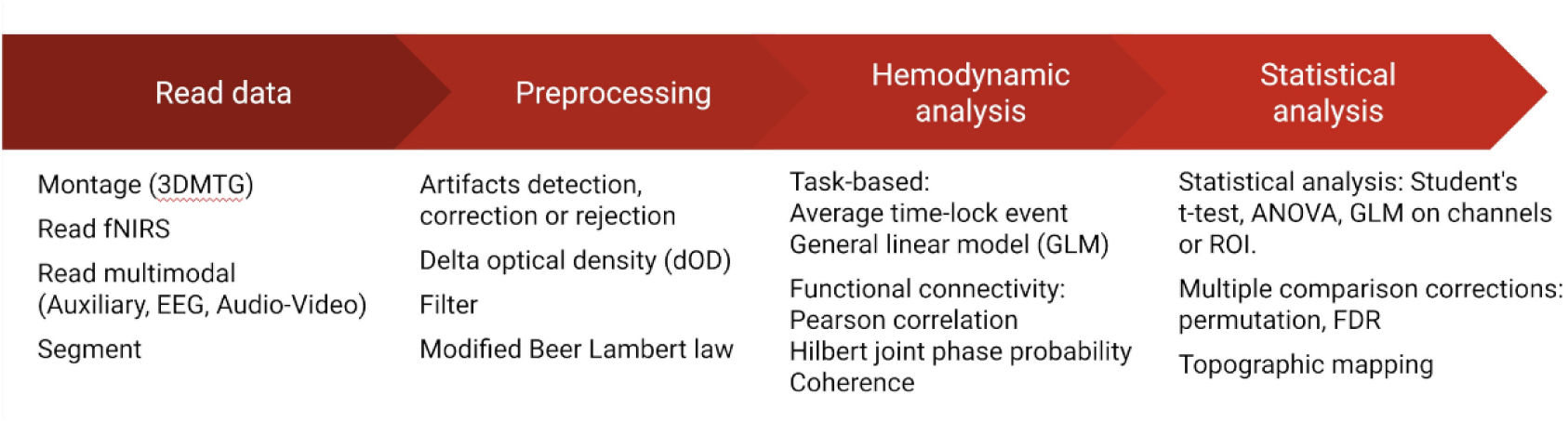
Overview of the organization and functionalities of the LIONirs toolbox.

Hemodynamic analysis involves the identification of the task-related hemodynamic response (HRF), by averaging the time-lock events or estimating of the relevant activities using the HRF model in a GLM. Functional network organization can be estimated using several brain functional connectivity (FC) measures such as Pearson correlation, Hilbert joint phase probability distribution and magnitude squared coherence (Xu et al., 2015; Molavi et al., 2014; Kida et al., 2016). Finally, some basic statistical analyses (e.g. student t-tests, ANOVA, GLM) can be applied to task-based components or FC measures. A correction for multiple comparisons using false discovery rate adjustment or permutation tests is also implemented. These results can then be displayed as a topographical representation of all channels, or a selection of them.

In the next sections (Sections 2.2 to 2.8), the main analysis steps and functionalities will be explained in detail and illustrated with examples. Several tutorials are available in the toolbox documentation, offering a step-by-step guide for some processing tools, along with data examples (https://github.com/JulieTremblay3/LIONirs).

### 2.2. Montage configuration: 3DMTG interface

In fNIRS, the position of each source and detector (optodes), commonly called the montage, is not standardized and must be adjusted to fulfill the experimental requirements. It is up to the user to associate sources and detectors with 3D coordinates, according to the specific montage of their study. Therefore, depending on the number of optodes available, the type of caps that is used, and the head sizes of the participants, the configuration is optimized in order to cover the regions of interest (ROIs). When determining a montage, it is important to match brain anatomy with the position of the optodes, in order to obtain reliable topographical and cortical projections. The optodes’ locations on the head of a subject are recorded using a stereotactic system or a 3D localization system, and saved in a coordinates file (.elp file). The 3DMTG interface reads the coordinates file, allowing the visualization of the optodes’ registered position on the 3D representation of the scalp or cortical surface (Figure 2). The correspondence between optode location and anatomical representation is done with a rigid body transform method, or on a real subject using anatomical markers (nasion, inion, left and right pre-auricular points) identified through structural MRI images (Penny et al., 2011). The anatomy can be defined by either an MRI template or the subject’s images (Richards et al., 2016). Optionally, MRI images include anatomical atlases such as a segmentation according to the Brodmann atlas or the automated anatomical labeling atlas (AAL), used to label functional cortical regions over the surface of the brain (Aleman-Gomez et al., 2006; Talairach, 1988; Tzourio-Mazoyer et al., 2002). The visualization of the atlas helps the user ensure that the montage adequately covers the ROI of the cerebral cortex. If the individual’s MRI images are available, the anatomy of the brain and skin surface can be extracted using an external image processing software such as Neuronic Image Processor (http://www.neuronicsa.com/modulos/producto/imagic.htm) or Brainsuite (Dogdas et al., 2005). The toolbox also provides a generic template in the required format for projection onto the skin and cortical surface (Collins et al, 1994). When no MRI data is available for an individual, the brain MRI template will likely require spatial rotation, translation, and scaling adjustments, in order to match the individual’s optodes coordinates. Once all these steps are completed, the experimental project file can be saved. This allows the user to access the topographical representation of the data at any stage of the analysis and visualize the entire or partial data on the skin or cortex.

**Figure 2.**
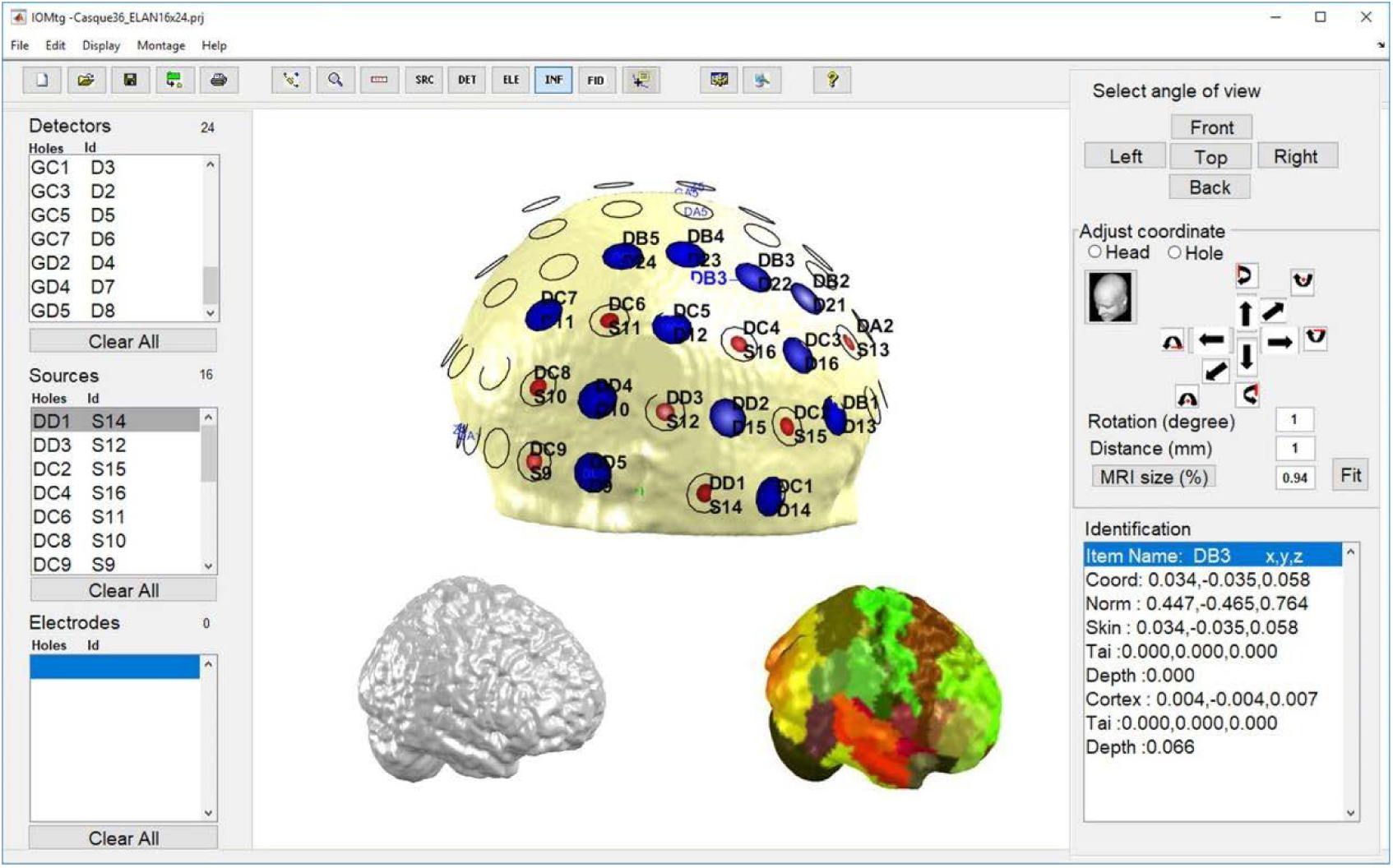
3DMTG interface for creating and visualizing the fNIRS optode localization. The position of sources (S) and detectors (D) can be projected onto the head surface. The skin (top image), cortical surface (bottom left image), or associated anatomical regions (bottom right image) can be used as anatomical landmarks to dictate the placement of the helmet holes. Each hole has a label that allows the identification of the physical holes on the cap. The three panels at the left list the associations between the labels of detectors (top), sources (middle) and electrodes (for simultaneous NIRS-EEG recordings; bottom) with the helmet holes. The upper panel contains tools for moving the entire head, adding or removing sources, detectors and electrodes, and accessing parameter adjustments. The right panels allow the user to select the angle of visualization (top), adjust the coordinates of the registered positions of hole(s) (middle) and provides information about exact coordinates (bottom).

### 2.3. Data organization

LIONirs supports many raw data formats, including ISS, NIRx, .nirs, and SNIRF. It reads the data and uses the previously created 3DMTG anatomy to attribute each channel to the correct location on the scalp. Simultaneously acquired multimodal data, such as physiology, EEG, or audio-video recordings, can be integrated and synchronized, becoming accessible through the DisplayGUI. A trigger sent simultaneously to all equipment (e.g. fNIRS system and EEG amplifier) is essential for the proper synchronization of multimodal data. Any step of processing and adjustment taken in LIONirs can be saved to different folders, organized in a tree structure for each subject, as depicted in Figure 3. The necessary data and associated Matlab structure (NIRS.mat) are saved to each (sub)folder corresponding to the various stages of processing. This NIRS.mat structure allows one to easily keep track of all processed operations and apply modifications on one level, without having to start from the very beginning. Folders and subfolders are also used to separate branches of a processing pipeline for parallel analysis of multiple conditions or different analytical approaches. Finally, at every stage, the data can be exported to the most common fNIRS format (.nirs or SNIRF), thereby facilitating compatibility with other software.

**Figure 3.**
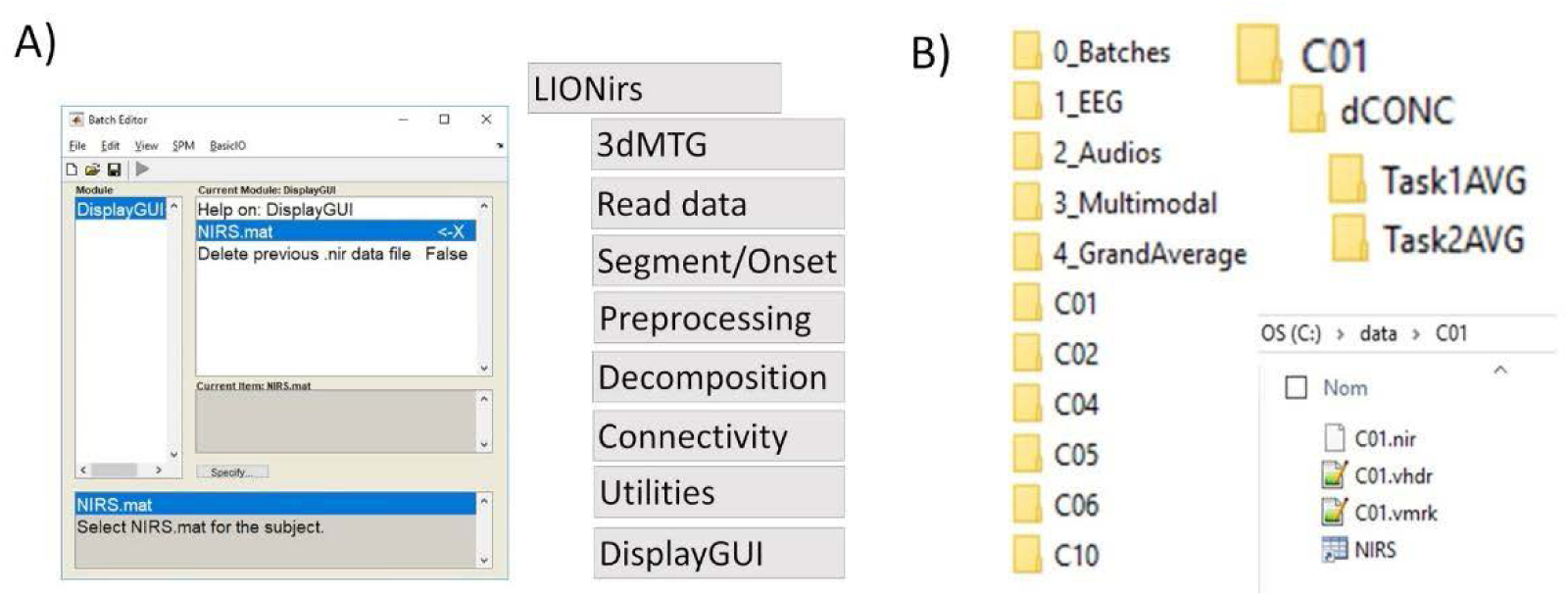
Organization of the different stages of data analysis. A) The Matlab batch editor gives access to the LIONirs functions menu (e.g. artifact detection, filtering, averaging, etc.), allowing the user to build up a processing pipeline for their specific needs. B) Example of folder and subfolder organization of the various processing steps applied at the subject or group-level. Each folder contains the NIRS.mat structure, to keep track of the operations applied and the analyzed data files. Subfolders can include separate branches for different parallel pipelines, for example, when analyzing multiple experimental conditions.

### 2.4. Visualization of the data (DisplayGUI)

One of the main objectives of this toolbox is to offer as much flexibility and transparency as possible, so that the user need not have programming skills in order to interact with the data. For this purpose, the DisplayGUI interface allows one to visualize the data at any step of the analysis. Figure 4 shows an overview of the DisplayGUI, including the multimodal display, different decomposition techniques, and the 3DMTG navigation. After synchronization with fNIRS, multimodal data such as respiratory signal, cardiac pulse, or other auxiliary channels are displayed in the upper window of the DisplayGUI. Video, EEG or 3DMTG topographic representation can be displayed on demand for a selected time interval. Specific spatial ROI can be selected in the 3DMTG interface to restrict visualization in the DisplayGUI and, inversely, channels of interest can be selected in the DisplayGUI, leading to a restricted visualization in the 3DMTG.

**Figure 4.**
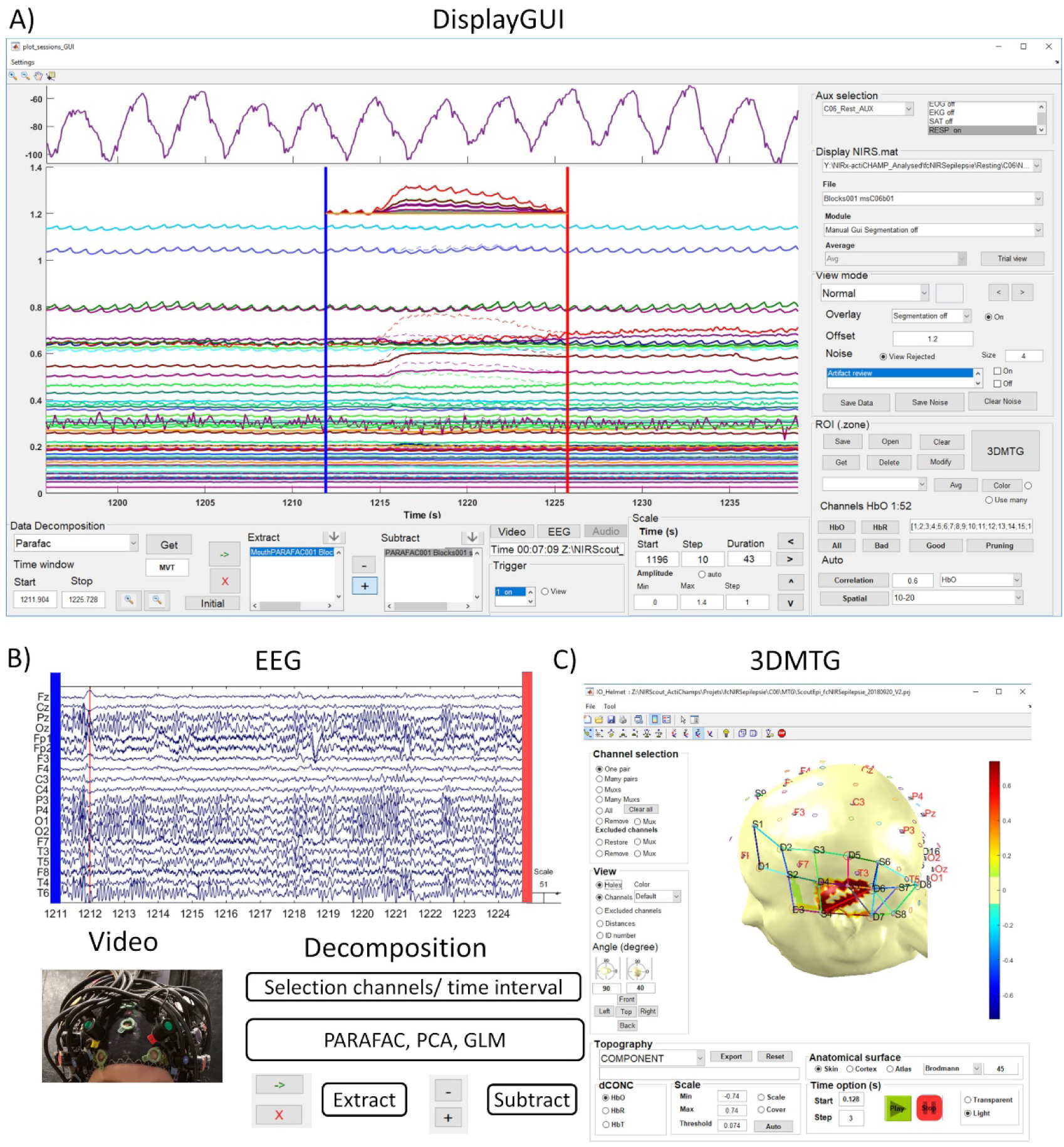
Overview of the DisplayGUI interface for visualization of data at each processing stage. A) The upper panel shows the various multimodal recordings that can be synchronized with the fNIRS data, e.g. auxiliary for respiratory belt trace (purple line). The large panel below shows the fNIRS data at a specific stage, with different colors for each channel. Within this large panel, the right column includes different functions allowing the user to choose which fNIRS signal to display [Display NIRS mat], navigate between different blocks [File] and processing steps [Module], options to visualize channels using a butterfly or a standard view [View mode], and overlay the signal of a previous step with the current module [Overlay]. The user can also define [Get] and save a list [Save] of regions of interest [ROI zone], and use these to display a subset of channels. In the lower part of the large fNIRS data panel, one can access different data decomposition techniques, such as GLM, PARAFAC, and PCA, in order to identify components corresponding to artifacts or relevant physiological activities. The component of interest can subsequently be extracted for further analysis or, in case of artifacts, be subtracted from the data. Finally, the lower part of the panel at the right provides options to display [Trigger] or adjust [Scale]. B) Using the time start (blue vertical line) and time stop (red vertical line) options on the displayed fNIRS data (in A), one can select a window of time for which to quickly access the simultaneous EEG, audio-video recording, reject a section of data or apply a decomposition technique. (C) Topographic visualization of the fNIRS signal using the [3DMTG]. The channels and hemodynamic activity can be projected onto a 3D representation of the skin (as shown on the Figure), the helmet or the cortical surface.

The user can either look at individual channels, select a group of channels based on the correlation of their time courses, or choose channels based on their amplitude or spatial localization. Subsequently, the user selects a specific time interval, such as the duration of an artifact or a task. Specific processing steps, such as rejection of noisy intervals or correction of artifacts, can be performed for the selected time interval and specific channels. Various data decomposition techniques are implemented in such a way that the user has the possibility to first extract one or several specific components, which will then be stored in the component list [Extract]. Because many of the decompositions can be influenced by the presence of outliers in the data, the customized time and channel selection within the DisplayGUI is a particular advantage of the LIONirs toolbox, favoring a superior and more targeted decomposition. Visualization of the components’ signatures in all considered dimensions, i.e. time, space, or wavelength, allows for easy identification of the components of interest. For instance, when aiming to correct an artifact, the user can identify those specific components with characteristics related to that artifact, and subtract it from the data. When decomposing a cerebral activation pattern, the visualization of the components allows for easy identification of those components that contain characteristics known to be related to a hemodynamic response, and should therefore be exported for statistical analysis (section 2.8). If one has reason to exclude some or all channels of a specific interval, for example due to noise contamination, the DisplayGUI allows a manual rejection to be applied. Even though the DisplayGUI can visualize data from previous processing steps, the operations are done in a sequential manner. This means that any manual editing (e.g. artifact correction) is applied to the latest data stored to memory. However, the user has the option to overlay the data from the previous step onto the signal of the current module, to fully appreciate the effect of the procedure. This is, for example, useful during artifact correction, when overlaying the original onto the corrected one provides a better picture of the correction’s effect on improving the signal’s quality. If the corrections do not lead to satisfying results, it is possible to restore the previous data in the DisplayGUI.

### 2.5 Preprocessing tools

#### 2.5.1. Channel quality check

In fNIRS data analysis, a common way to ensure signal quality is to detect the cardiac pulse, which is an easy-to-monitor physiological marker (Fekete et al., 2011; Goodwin et al., 2014; Pinti et al., 2019). Hence, in the absence of a cardiac pulse due to poor contact of the optodes with the skin, specific channels should be excluded. The frequency of the cardiac pulse varies with age, from 2 Hz at birth (Mortensen et al., 2017) to around 1 Hz in healthy adults (Fekete et al., 2011). In the software implementation, the user can adjust the sensitivity criteria in order to discard those channels that are without evidence of cardiac pulse, and thus ensure the quality of the fNIRS data. The toolbox offers the possibility to save a report that provides information about the peak frequency identified and which channels were excluded.

#### 2.5.2. Automated artifact detection and correction

Abrupt variations in the raw fNIRS signal due to motion or muscular artifacts are often easy to detect (Schecklmann et al., 2017; Scholkmann and Wolf, 2013). The batch system has an artifact detection module that can automatically identify intervals and channels affected by a suspected artifact. The detection is performed using three criteria, whose sensitivity can be fine-tuned by the user. A first criterion uses a moving average technique to detect abrupt variations of amplitude that are abnormal; those intervals are identified when the variation is higher than a predefined normalized z-score threshold, computed on the channel’s whole recording (Weinberg and Abramowitz, 2008). This method avoids over-sensitivity to background noise and, more specifically, detects excessively high variations of amplitude. A second criterion marks any artifact-free data segment that is between two artifacted segments, and shorter than a minimal time interval specified by the user. A third criterion uses the temporal correlation between channels with artifacted segments (detected using the two previous criteria) and any other channels. This implies that if the correlation between the time courses of a channel that contains an artifacted segment and another channel is higher than the threshold defined by the user, then the time interval of this other channel is also identified as containing an artifact. It is then up to the user to conduct a manual verification of the artifact detection and potentially apply adjustments to either reject the identified segments, or to proceed to one of the artifact-correction techniques provided in the toolbox. These corrections aim to minimize the overall effect of artifacts on the fNIRS signal, by avoiding the rejection of too many trials and thus maintaining adequate statistical power for further analyses.

LIONirs toolbox offers two data decomposition techniques specifically for artifact correction: the 2D PCA and the multidimensional PARAFAC. For both approaches, LIONirs allows the user to conduct an automated decomposition of segments previously identified as containing artifacts. Both have been proven to perform well for correcting movement artifacts in fNIRS (Hüsser et al., 2019; Yücel et al., 2014). The implementation of PCA uses a Matlab function to decompose time and channel into components that represent most of the variance, and classify these by percent of variability explained. With PCA, we assume that the component that explains most of the variance will reflect the artifact, and the user can remove the component up to a total percentage of variance explained. (Cooper et al., 2012; Brigadoi et al., 2014). The implementation of PARAFAC is done by means of the external open-source package Nway Toolbox (Andersson and Bro, 2000). In PARAFAC, an automated decomposition selects the optimal number of components, based on the lowest residual error and the highest Concordia criteria (Andersson and Bro, 2000). The artifact component(s) retained for correction are the one(s) that explain the highest variance in the time course (similar to PCA) and also the lowest difference between wavelength, assuming that both wavelengths from the same site will perceive the artifact conjointly (Cui et al., 2010).

For both decomposition techniques, the user can choose between an automated subtraction using the batch system or a manual one using the DisplayGUI. Once the decomposition completed, the final effect of the correction can be reviewed visually in the DisplayGUI. There is also an offset adjustment available to force light intensity to return to the level it was at before the movement artifact, and thus avoid discontinuity in the signal. This procedure greatly improves data quality, as illustrated in Figure 5. In this case, the uncorrected artifact shows both wavelengths equally affected by an articulatory movement and an abnormal strong variation, while the corrected one shows the slow pattern of the HRF time-lock with the task event.

**Figure 5.**
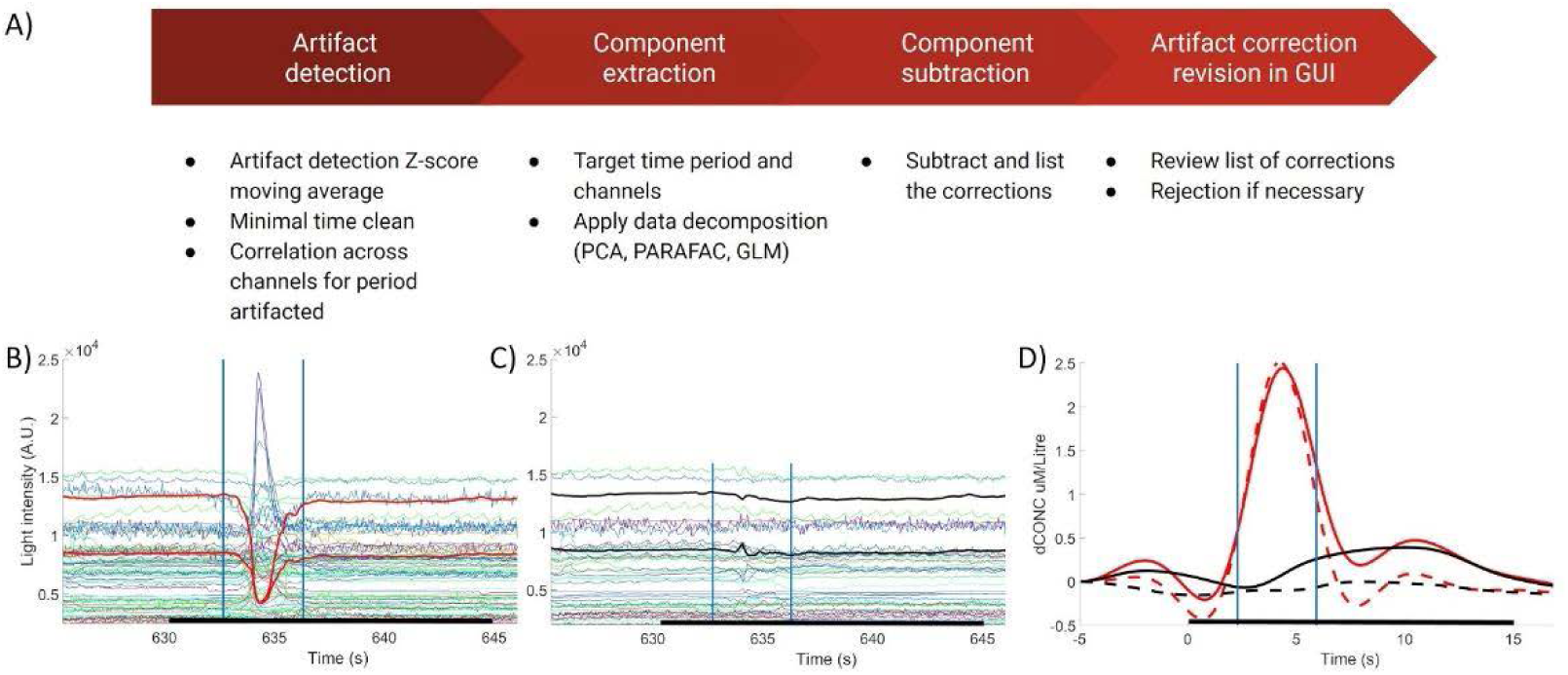
Example of a movement artifact correction. A) The proposed sequence for the automated detection and correction of artifacts in the LIONirs toolbox. B) 15-s segment (black horizontal line) of raw fNIRS data recorded during a verbal fluency task, where the participant has to say out loud as many words as possible belonging to a given category during a specific time interval (Paquette et al., 2015). This task typically induces an articulatory movement artifact, expressed as an abrupt change in the fNIRS signal (shown on B between the two vertical blue lines). C) The same data segment after the subtraction of the artifact-related component, identified with PARAFAC decomposition. D) Concentration changes in HbO (solid lines) and HbR (dotted lines) from the channel which most prominently showed the artifact in the raw data, before correction (red lines) and after correction (black lines). HbO and HbR signals are both equally affected by the artifact (red lines) as they both show a strong abnormal variation, while the corrected data (black lines) reveals the slow pattern of the HRF time-lock with the task event in the Broca ROI typically recruited by the verbal fluency task.

Although this article has laid out the different options for detecting, correcting and rejecting artifacts in the order in which they are typically performed for movement artifact correction, the LIONirs toolbox offers the flexibility to change that order, and to apply each method at any stage of processing.

#### 2.5.3. Systemic physiology artifacts

fNIRS measures light attenuation as deep as a few centimeters under the surface of the skin (Haeussinger et al., 2011). Light absorption, however, is not specific to the cortex, as photons travel through skin, skull, and other superficial tissues. Thus, systemic hemodynamic variations related to other physiological phenomena not directly related to cerebral activation, such as cardiac pulse, respiration, or slow-wave fluctuations around 0.1 Hz also known as Mayer waves (Julien, 2006), are part of the acquired signal and distort the hemodynamic response. Signals related to cardiac pulse or respiration can be filtered out, as their frequencies are not in the range of the hemodynamic activity (Fekete et al., 2011).

However, systemic slow hemodynamic fluctuations such as Mayer waves could be confounded with the hemodynamic response associated with brain activity. Multi-distance optodes, global average of all channels or PCA of the entire signal that includes the slow systemic fluctuations, can be used to identify and correct this type of noise (Erdoğan et al., 2016; Gagnon et al., 2012; Goodwin et al., 2014; Kirilina et al., 2012; Saager and Berger, 2005; Zhang et al., 2005). The GLM (see section 2.6.1.) can be applied, to estimate the regression coefficient corresponding to the slow systemic physiological noise, which can then be subtracted from the data. Regressing the confounding physiology increases the sensitivity of the subsequent analysis, thus improving the reliability of the results (Zhang et al., 2005; Huppert, 2016). Figure 6 illustrates an example of such a regression using globally averaged data (all channels averaged) to estimate the effect of physiology artifact as a regressor (Figure 6B). The HRF response is more stable after (Figure 6C), compared to before (Figure 6A) the extraction of the physiology.

**Figure 6.**
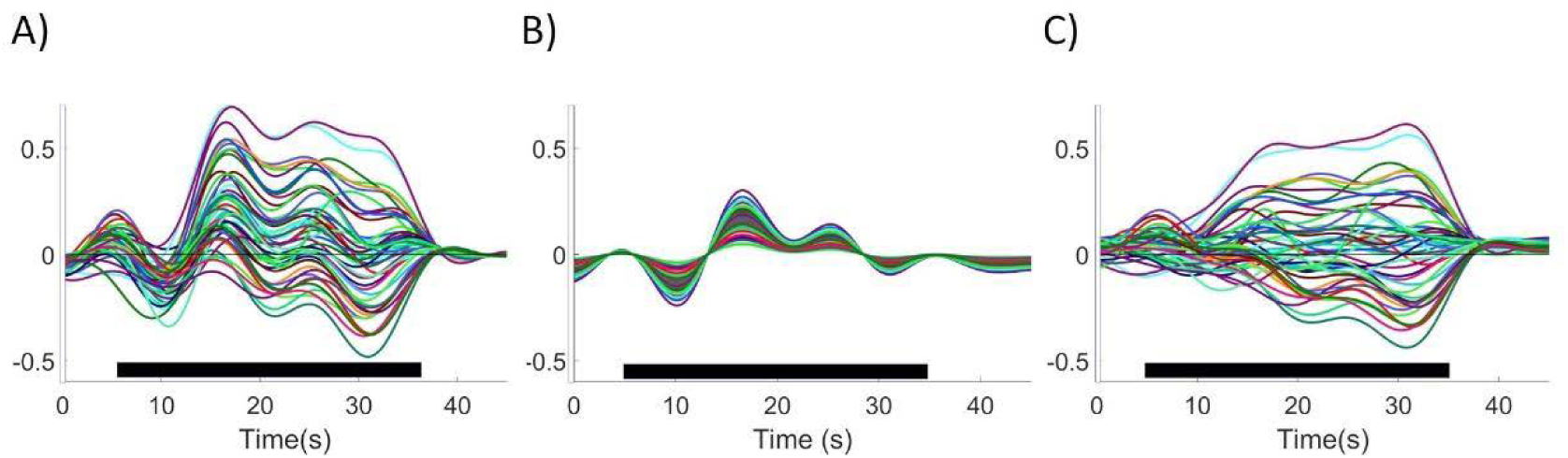
Regression of the physiology, applied to a single trial of a fNIRS data set recorded during a 30-seconds visual stimulation (black horizontal bold line) with a flipping checkerboard. A) The raw data from the 54 fNIRS channels covering the occipital area; B) The extracted physiology artifact using a GLM regression in the same fNIRS channels. Slow-wave fluctuations around 0.1 Hz are clearly visible in the physiology extraction. C) fNIRS data after subtraction of the physiology artifact.

### 2.6. Hemodynamic analyses

Once data is corrected, by subtracting or rejecting confounding artifacts, the task-related brain activity or functional connectivity (FC) measures can be identified for each participant. LIONirs preprocessing modules handle the basic transformation of the data, such as delta optical density (dOD), filtering, and transformation of light intensity into hemodynamic concentrations using the Modified Beer-Lambert law (MBLL) (Delpy et al., 1988). The differential pathlength factor (DPF) can be adjusted according to the participant’s age (Duncan et al., 1996, 1995; Scholkmann and Wolf, 2013) or be fixed manually. When the signal has been transformed into hemodynamic fluctuations, there are different ways of proceeding to the interpretation of cerebral activation. For example, it is possible to process task-based data in order to identify the specific characteristics of the task-related hemodynamic response. Functional connectivity data acquired at rest or during a specific task can also be processed in order to investigate brain network organization. In the next two sections, we will present task-related and functional connectivity analyses, respectively, using the LIONirs toolbox.

#### 2.6.1. Task-related hemodynamic response

A typical way to estimate the HRF response is through a regression approach, namely the GLM, that allows the researcher to determine how well the data points match a set of predictions (Friston et al., 1995; Penny et al., 2011; Poline and Brett, 2012). A multiple linear regression based on the least-squares estimation has been implemented into the current toolbox, where the user must define the regressors to be considered (Draper, 1998). A tool is provided to convolve the modeled HRF based on the applied paradigm (Glover, 1999) and to create an event-related regressor. To improve performance of the GLM, the user has the flexibility to add relevant regressors to the model, i.e. factors that might be correlated with each other, such as auxiliary data (e.g. cardiac fluctuations or respiration) (Hillenbrand et al., 2016) or low-frequency oscillations (Tong and Frederick, 2010). For example, one or several short-distance channels (Gagnon et al., 2012), or the global average (Zhang et al., 2005) can be used to estimate the physiological noise. Once the GLM estimation of the HRF response is obtained at the subject-level, individual regression weights, typically called beta coefficients, can be exported for use in group statistics. Decomposition using PCA or PARAFAC to extract the hemodynamic time course, could be explored as well. As a visual example, Figure 7 shows a typical processing pipeline (7A) for data acquired during a visual stimulation (7B) and a data set acquired during a passive story listening paradigm (7C). Data were recorded using an Imagent Oxymeter (ISS, Champaign, Illinois, USA) on a healthy adult subject, at a sampling rate of 20 Hz. Experiments were approved by the ethical committee at the Sainte-Justine University Hospital. The following modules were applied to the data (see also Figure 7A): read data, add multimodal data (e.g. HRF model response, video), segmentation to synchronize multimodal data, semi-automated artifact detection, and rejection, dOD, low-pass filtering at 0.1 Hz, and MBLL. Then, a GLM model, including global average (Zhang et al., 2005) and the modeled HRF as regressors (Glover, 1999), was applied. The estimated physiology artifact was subtracted from the data, before averaging across trials to obtain the time-locked response. The estimation of the HRF response was followed by a simple Student t-test against zero, applied to all coefficients that were estimated from multiple sessions for each channel (Uga et al., 2014). False discovery rate (FDR) was used to correct for multiple comparisons (Benjamini and Yekutieli, 2001). The response that differed significantly from zero was projected onto the cortical surface.

**Figure 7.**
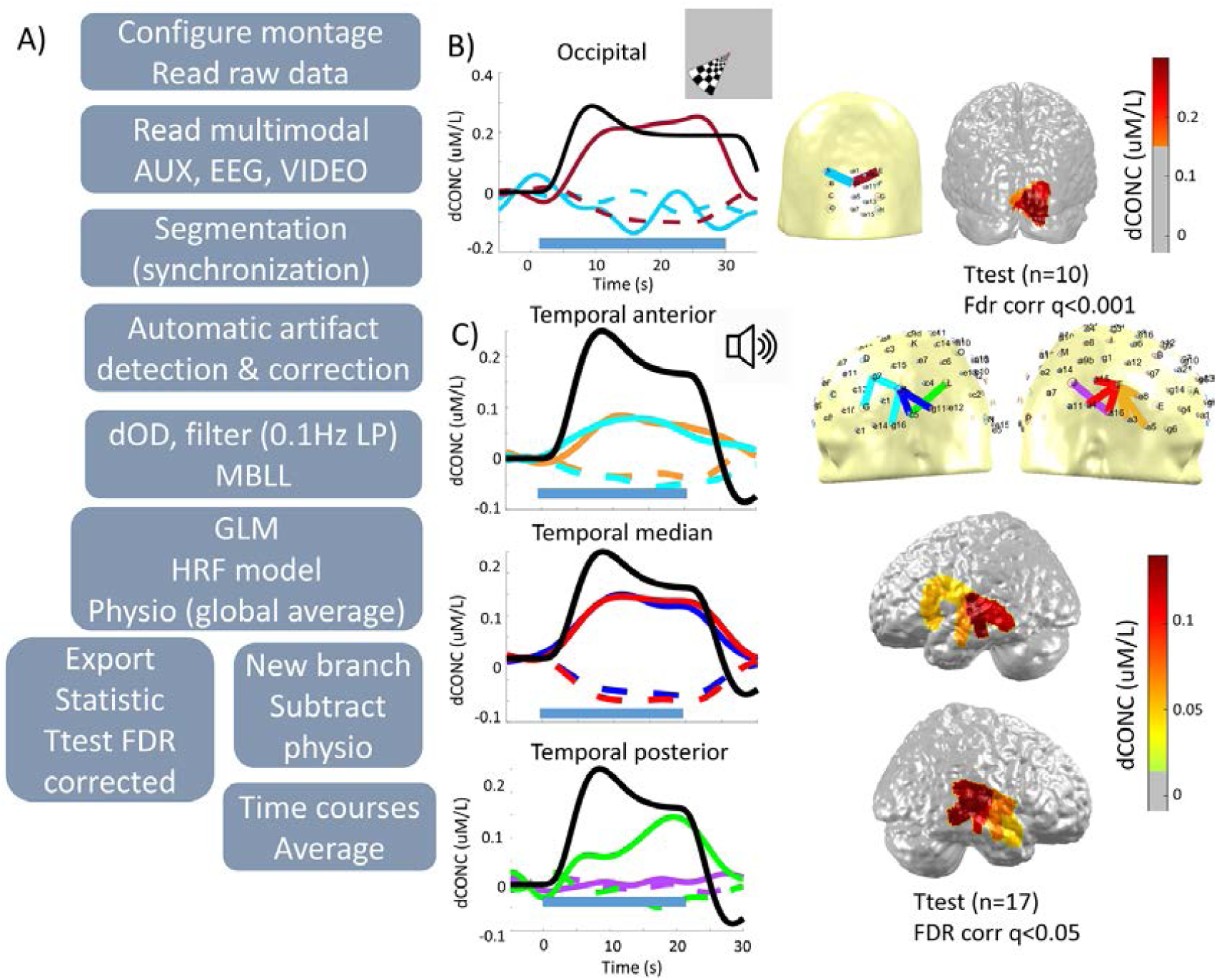
Examples of subject-level analyses. In both cases, the results concord with previous fMRI analysis of similar experiments (Toronov et al., 2007; Vannest et al., 2009). A) Detailed hierarchical diagram of the different steps included in the processing pipeline. B) **Visual stimulation**. Fifty-four fNIRS channels covered the occipital cortex, measuring the cerebral activation in a healthy adult. Visual stimulation consisted of a flipping checkerboard that appeared for 30 seconds in the bottom-left corner of the stimulation screen (Bastien et al., 2012). The task included a total of 10 blocks. Averaged time courses of all blocks are shown by a solid line for HbO and a dotted line for HbR concentrations. The color of each curve associates with a channel(s) at a specific location on the head topography. The duration of stimulation for each paradigm is shown with a horizontal blue line below the timeline curve. The black line displays the expected HRF model. The significant response for HbO variations were projected onto the cortical surface (right panel). A strong statistical response was found in the right visual cortex (FDR corrected q<0.001). C) **Passive story listening task**. One hundred ten channels covered the bilateral frontal, temporal and parietal areas. Hemodynamic fluctuations were recorded while the subject listened to a story presented in their native language. The story was presented in 18 individual blocks of 20 seconds each (Paquette et al., 2010). Note that the description used in B is also used here to illustrate HbO and HbR concentrations (solid and dotted lines, respectively), the duration of the task (horizontal blue line) and the expected HRF model (black line). Temporal anterior and temporal medial areas of both hemispheres showed significant responses related to the auditory response, while the temporal posterior area in the left hemisphere revealed cerebral activation most likely related to receptive language processing in Wernicke’s area (FDR corrected q<0.05).

#### 2.6.2. Functional connectivity measures

Functional connectivity (FC) is usually defined as the correlation between the time series of hemodynamic fluctuations measured at different locations on the scalp (Friston, 2011). FC has been largely studied in fMRI, and assumes that there is a direct relationship between neuronal activity and the hemodynamic fluctuations, measured as the BOLD signal. Research in this field has allowed the description of several consistent functional brain networks during resting state, including the default mode network, the somatosensory network, the dorsal attention network, the ventral attention network, and the auditory and visual networks (Bellec et al., 2010; Damoiseaux et al., 2006; Bijsterbosch et al., 2017; Raichle et al., 2001).

Many of the mathematical measures that are used to estimate FC from neuroimaging time series are also frequently applied in other modalities, such as fMRI or EEG (Friston, 2011; Smitha et al., 2017). Most of these approaches can be applied straightforwardly to the time series of fNIRS data (Gallagher et al., 2016; Molavi et al., 2014; Nguyen et al., 2018; Xu et al., 2015). However, fNIRS is limited to measuring hemodynamic fluctuations that occur in the cortical surface, thus precluding fNIRS data from informing about functional brain networks in deeper areas. Importantly, depending on the level of coverage of the montage used and duration of the recording, connectivity measures derived from fNIRS data should be interpreted carefully (Sun et al., 2018; Wang et al., 2017). Despite these limitations, FC measures can be applied to fNIRS data in order to answer specific questions regarding the characteristics of cortical functional networks in a specific population and/or condition.

The LIONirs toolbox allows the computation of functional brain connectivity measures within the predefined channels or ROIs, using Pearson’s correlations (Xu et al., 2015), Hilbert joint phase probability (Molavi et al., 2014), and magnitude squared coherence (Kida et al., 2016). Figure 8A shows a typical processing pipeline for performing a FC analysis. In this example, we applied the magnitude squared coherence approach to fNIRS data recorded in 14 healthy adults during a 12-minutes resting-state period. Based on the 10-20 system (Klem et al., 1999) several ROIs were created in the DisplayGUI by assigning each channel to a specific ROI (Figure 8B). For each participant, magnitude squared coherence measures are computed between the time series of each pair of channels and are presented by ROIs in a FC matrix (Figure 8C) or a circular connectogram (Figure 8D) using LIONirs GUI_LookMatrices and user-defined thresholds. FC measures could also be computed between each group of channel(s) representing ROIs (not illustrated in Figure 8). The results of the resting-state coherence matrix for all healthy adults include the averaged coherence values for all subjects. Some connectivity clusters are observed in the prefrontal areas (FP1 and FP2), the somatosensory areas (F3, C3 and F4, C4) as well as the left-hemisphere language areas (F7 and T3, T5).

**Figure 8.**
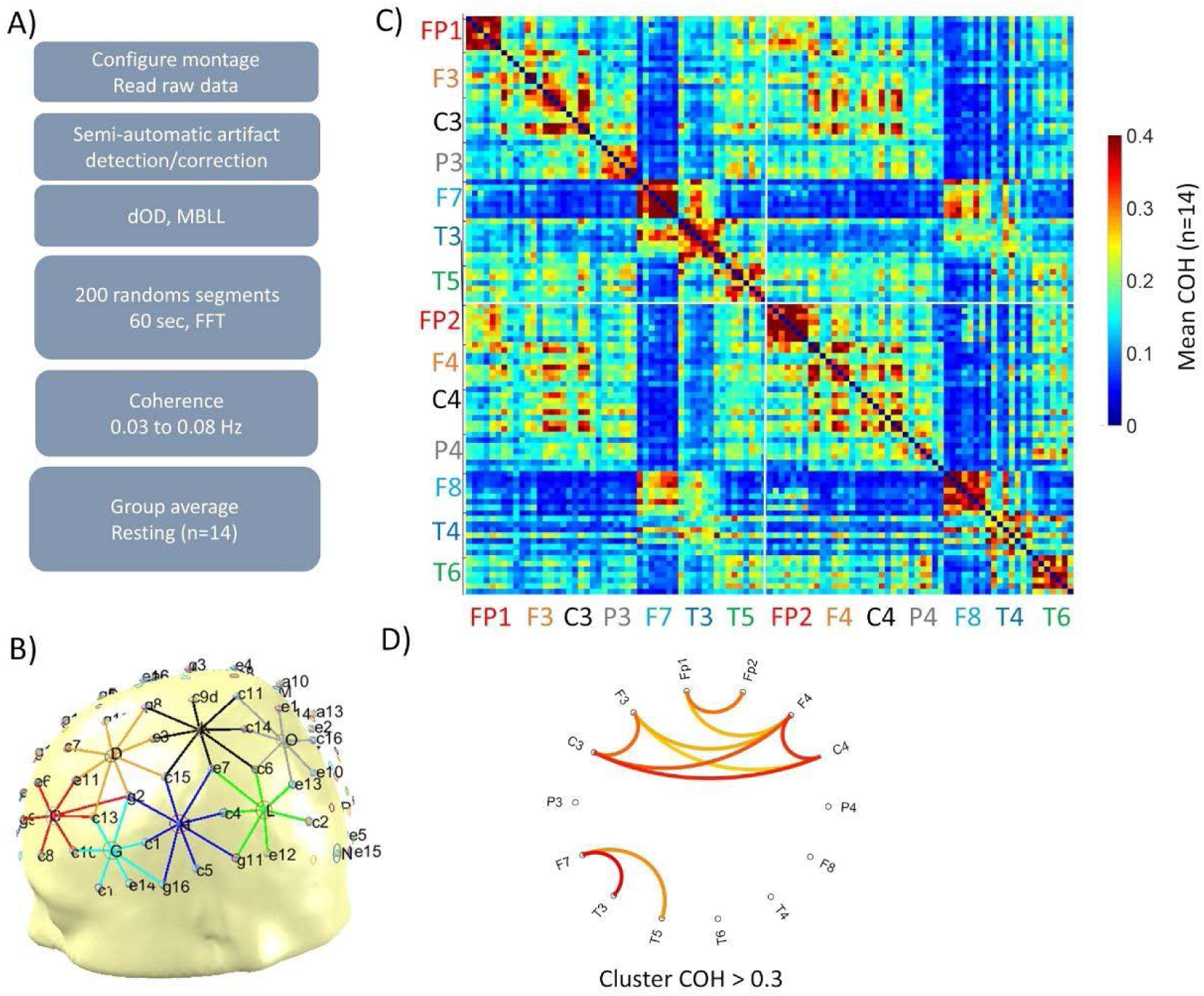
Functional connectivity (FC) analysis for a fNIRS data set of 14 healthy adults during a resting-state data acquisition (eyes open, 12 minutes). A) A typical processing pipeline used to calculate FC using magnitude squared coherence of HbO hemodynamic concentrations. B) Regions of interest (ROIs) were defined manually using the DisplayGUI (the montage was identical for both hemispheres and the same for all participants). These ROIs were used to compute the connectivity matrix (shown in C). Red channels (B and C) are located around Fp1 and Fp2 from the 10-20 system; orange channels around F3 and F4; black channels around C3 and C4; gray channels around P3 and P4; cyan channels around Fp7 and Fp8; blue channels around T3 and T4; and green channels around T5 and T6. C) A coherence matrix between all pairs of ROIs is averaged for all 14 subjects (average matrix shown in this figure). In this matrix, the upper left quadrant corresponds to ROIs in the left hemisphere, the lower right part corresponds to the right hemisphere, while the lower left and upper right quadrants reflect inter-hemispheric connections. The color scale ranges from blue to red, where blue means there is no or weak functional connectivity and red suggests highly functionally connected regions. D) The connectogram reveals the stronger connections (COH>0.3) across the ROIs.

### 2.8. Statistical analysis

Statistical tools included in the LIONirs toolbox can be applied to specific channels or ROIs. Hence, prior to the analysis, it is essential that the user identifies homologous channels or ROIs for all subjects, to allow for proper inter-subject comparisons. Statistics can be applied to the averaged time course, to the components identified with a decomposition technique (e.g. the estimation of the hemodynamic response with the GLM model), or to FC measures. The statistical tools integrated in the LIONirs toolbox are the parametric student t-test and analyses of variance (ANOVA). A FDR correction for multiple comparisons (Benjamini and Yekutieli, 2001) is also available. Permutation tests have been implemented for parametric student t-test, which establishes the null distribution based on experimental data shuffle test repeated multiple times (Galán et al., 1997). The results can be easily visualized as a topographical projection onto the scalp or cortical surface using 3DMTG. Even though statistical methods are supported in the current toolbox, specialized statistic softwares such as the Statistical Package for the Social Sciences (SPSS) or R (www.r-project.org) are recommended for advanced analysis.

## 3. Discussion

LIONirs is a new open-source toolbox for the analysis of fNIRS data. The toolbox includes a variety of techniques and methods for data processing that have been used for fNIRS data analysis in many previous studies. Since there is currently no consensus on the standard procedures for fNIRS data recording, preprocessing and processing, one of the main objectives of this work was to design a toolbox that is as flexible as possible, and allows for easy handling of the data without requiring programming skills. Hence, the LIONirs toolbox makes it possible for the user to choose between several tools, explore and compare various methodological approaches, and easily build a customized data analysis pipeline adapted to their specific needs and data set characteristics. The pipeline can subsequently be applied in either a fully or semi-automated manner to a large number of subjects. The second goal of the LIONirs toolbox was to provide an open-access and transparent tool for fNIRS data processing, thus avoiding the black-box phenomenon. To that end, two graphical interfaces (DisplayGUI and 3DMTG) provide 3D visualization of the fNIRS montage and the data at any stage of processing. It is thus possible to track each intermediate result across the entire data analysis process, allowing the user to modify, explore and compare the applied methods if necessary. fNIRS data acquired in children or clinical populations often includes a large number of movement artifacts that can rarely be fully corrected using an entirely automated procedure. A careful artifact detection and correction is critical for obtaining an interpretable fNIRS signal. The LIONirs toolbox enables a semi-automated artifact detection, correction, and rejection, allows the user to verify the quality of the preprocessing, make adjustments if needed, and combine different methods when applicable. LIONirs includes two data decomposition techniques for artifact correction, namely tPCA and PARAFAC, which are powerful tools for minimizing the impact of artifacts and increasing the quality of data (Yücel et al., 2014; Hüsser et al., 2019). A quality report can be produced to summarize the number and duration of the corrections applied for each subject. In some cases, poor signal quality may require the rejection of a channel or time interval instead of trying to correct it; an advantage of LIONirs is that it offers the user the flexibility to choose between either correction or rejection. Figure 5 illustrates the impact of non-corrected artifacts on the hemodynamic response, which could lead to misinterpretation of the data. The detection and correction or the rejection of artifacts are therefore crucial steps in fNIRS data analyses.

Another type of noise that often affects hemodynamic signals are slow-waves related to systemic physiological fluctuations (Yücel et al., 2016; Chaddad et al., 2013; Masataka et al., 2016). Several studies have shown that removing these slow-waves leads to a significant increase of the specificity of the fNIRS signal (Erdoğan et al., 2016; Gagnon et al., 2012; Goodwin et al., 2014; Kirilina et al., 2012; Saager and Berger, 2005; Zhang et al., 2005; von Lühmann et al., 2020). This correction is even more important when the hemodynamic response shows small amplitudes, due to a low signal to noise ratio (Tachtsidis and Scholkmann, 2016). To deal with this physiological noise, the LIONirs toolbox offers a simple regression of either short-distance channels or a global average of all channels. We have illustrated how useful a GLM regression is for extracting physiological noise, thereby improving the interpretability of the hemodynamic response (Figure 6; Uga et al., 2014; Scholkmann et al., 2014; Peng et al., 2016).

Although the debate is ongoing as to which of the current methods is best, LIONirs incorporates some of the commonly applied techniques for analyzing hemodynamic response. They include epoch averaging (illustrated in Figure 7) and multiple regression, which are mainly intended for task-related data sets. Several measures of functional connectivity, which can be applied to both resting-state and task-related data, are also available in the toolbox, namely Pearson’s correlation, Hilbert joint probability distribution of the phase, and magnitude squared coherence (illustrated in Figure 8, Xu et al., 2015; Molavi et al., 2014; Kida et al., 2016). Measures can be calculated to obtain individual and then group functional connectivity matrices (Xu et al., 2015). The LIONirs toolbox therefore offers a variety of hemodynamic responses and functional connectivity measures, which can be selected according to the research question.

In some studies, fNIRS acquisition is conducted simultaneously with other modalities such as physiologic measures (e.g. cardiac pulse, respiration), EEG, or audio-video recording. The LIONirs toolbox allows the integration and synchronization of multimodal signals, which can increase the fNIRS signal quality, help with fNIRS data interpretation, and provide rich information on neurovascular coupling. For instance, physiologic measures can significantly help detect physiological artifacts and remove them from the fNIRS signal. The movements of the participant during the fNIRS recording can be captured on audio-video monitoring, which can contribute to the identification of movement artifacts. EEG recorded simultaneously with fNIRS may provide precious information on the participants’ state of consciousness (e.g. asleep, awake, drowsy or alert), or detect pathological alterations (e.g. epileptogenic activity) that can contaminate the fNIRS signal. Multimodal data acquisition is an important source of information for a better understanding of the neurovascular coupling (Lecrux et al., 2019). Extensive work on multimodal analysis in EEG and fMRI has shown that even if the electric signal from the EEG and the hemodynamic variations from the fMRI both evolve over a different time scale, subject-specific cerebral responses from both modalities can be useful for identifying relevant neuronal activity. Single-trial discrimination has also been used to construct EEG-derived fMRI activation maps (Philiastides and Heekeren, 2009), which could be applied to fNIRS. The concurrent analysis of these data can be helpful for the detection of artifacts and the control of data quality, as variations in experimental design has been shown to cause complexities in the hemodynamic response (Issard and Gervain, 2018). Thus, multimodal data can contribute to a better understanding of the relationship between the time series and the evolution across time of EEG rhythmic activity such as alpha, theta and gamma frequency bands, and has provided important insights into the characteristics of brain activity (Goldman et al. 2002; Martínez-Montes et al., 2004).

LIONirs can be viewed as a complementary tool to currently available software packages. As described in the introduction, other well-designed and highly useful tools such as HomER TM (Huppert et al., 2009), NIRS-SPM (Ye et al., 2009), Brain AnalyzIR (Santona et al., 2018) and FC-NIRS (Xu et al., 2015) have been publicly released in the last decades. While it also integrates some of the methods proposed by other toolboxes, LIONirs’ development was mainly driven by the specific needs of processing artifacted data acquired from challenging populations (very young children or clinical populations), which are the populations we typically recruit in our lab. In these contexts, the signal’s characteristics may be influenced by ongoing development or pathological phenomena, which have to be taken into account when processing and interpreting data. We therefore developed a highly flexible toolbox, allowing the user to apply various methods and visualize the data at any stage of the analysis. In addition, LIONirs includes PARAFAC, a new method for artifact correction and the extraction of relevant activities in fNIRS (Hüsser et al., 2019). Its integration into the Matlab Batch System, where processing pipelines can be customized, further contributes to this flexibility and enables the user to generate scripts that facilitate the automated analysis of large data sets. The DisplayGUI permits the visualization of data at any processing stage, thus contributing to both the transparency of the toolbox and to its flexibility, since alternative approaches can be explored and compared visually. Since montages are often not standardized across studies that use fNIRS, the 3DMTG GUI serves to create customized montages and associate them with a reconstruction of the brain’s anatomy, using the subject’s (or template) MRI images. Consequently, the user can, at any step during processing, organize the data by ROI. As do other fNIRS data processing toolboxes, LIONirs supports reading in and exporting to several data formats (e.g., .nirs, SNIRF, binary files), which allows the use of methods from other software packages.

As with other publicly available fNIRS data processing packages, LIONirs is in constant evolution, with new features and methods being continually added and improved. Novel functionalities will be progressively integrated into LIONirs and made publicly available. In the current version, the toolbox is limited to the visualization of the brain cortex by use of a simple radial projection from the corresponding channel position, because this provides an easy and fast (<1 sec) representation. Complete forward modeling and inverse estimation (diffusion optical tomography) are not yet supported, although they could be useful in studies that include high-density data acquisition. In a previous study, several methods were compared in a fNIRS context (Tremblay et al., 2018) and will be included in one of the upcoming versions of LIONirs. Nevertheless, the current version supports and provides several data formats; the integration of additional formats are planned, in order to make the toolbox available to a larger range of users. We also intend to integrate a technique for quantitative analysis of multimodal data, such as the N-way partial least-squares technique for a data-driven combination of neurophysiological EEG information and the hemodynamic response measured with fNIRS (Martínez-Montes et al., 2004).

Despite the examples used in this article to illustrate how a processing pipeline in LIONirs can be applied to real experimental data, we do not claim to establish a best-practice guideline for the analysis of fNIRS. The aim was rather to provide different options, so that researchers can build the processing pipeline that best suits their specific experiment and research question.

## 4. Conclusion

LIONirs is a new Matlab-integrated toolbox for fNIRS data analyses. It allows the user to deal with fNIRS data acquired with a wide range of optode montages, various correcting artifact methods, and multimodal data. The toolbox stands out because of its flexibility and transparency throughout the entirety of analysis processing. The user can explore and compare different methodological approaches and select the most appropriate technique for a particular data set. The toolbox provides the flexibility to build a customized processing pipeline, whereby the methods and order of analysis are completely determined by the user. Transparency is achieved via the DisplayGUI, that which allows one to visualize the continuous time series of the signal and a 3D projection of the data onto the scalp or cortex. The proposed semi-automated approach for artifact detection and data decomposition helps to efficiently reduce artifact contamination and promote quality of the fNIRS signal. LIONirs includes several techniques for the analysis of the hemodynamic signal, such as averaging, multiple regression, and FC measures, as well as a module for certain statistical analyses. The program is distributed under the GNU license and is open for further development and use in any third-party study, as long as this publication is cited. The current version, as well as detailed documentation on all its functionalities, can be downloaded from https://github.com/JulieTremblay3/LIONirs.

## Acknowledgments

The present work was driven by specific needs expressed by professors, students and collaborators working with fNIRS. Fruitful discussions and suggestions, ongoing for 10 years, have greatly improved the first core of methods for fNIRS data analysis. For that, we wish to thank Dr. Dang Nguyen, Dr. Frédéric Lesage, Dr. Maryse Lassonde, Dr. Franco Lepore, Dr. Renée Béland, Sébastien Bérubé, Hubert Jacob Banville, Mathieu Bissonnette, Olivia Florea, Danielle Bastien, Dr. Dima Safi, Melanie Lefrançois, Dr. Natacha Paquette, Solène Fourdain, Kassandra Roger and Justine Loignon-Lapointe.

## Funding

This work was supported by the National Science and Engineering Research Council of Canada (NSERC #2015-04199); the Canada Research Chairs (#950-232661), the Fonds de Recherche du Québec Santé (FRQS #28811) awarded to A.G.

## CRediT author statement

**Julie Tremblay:** Conceptualization, Methodology, Software, Writing-Original draft preparation, Validation

**Eduardo Martínez Montes:** Conceptualization, Methodology, Writing - Review & Editing

**Alejandra Hüsser:** Writing - Review & Editing, Resources, Validation

**Laura Caron-Desrochers:** Writing - Review & Editing, Resources, Validation

**Philippe Pouliot:** Conceptualization, Methodology

**Phetsamone Vannasing:** Conceptualization, Resources, Validation

**Anne Gallagher:** Supervision, Project administration, Writing - Review & Editing, Funding acquisition

## Declaration of competing interest

The authors declare no conflicts of interest with respect to the research, authorship, and/or publication of this article.

## Notes

https://github.com/JulieTremblay3/LIONirs

### Competing Interest Statement

The authors have declared no competing interest.

